# Small- and large-scale patches shape benthic microbial community structure and function in streams at the subcontinental scale

**DOI:** 10.1101/2025.10.08.681227

**Authors:** Richard LaBrie, Rebecca L. Maher, Kelly S. Aho, Brittni L. Bertolet, Nicholas E Ray, Paula C.J. Reis, Andrew L. Robison, Shannon L. Speir

## Abstract

Streams and rivers process dissolved and particulate matter as water moves along the land-to-ocean continuum, making important contributions to global biogeochemical cycles. Yet, predicting stream and river microbial metabolism associated with biogeochemical transformations at broad spatial scales remains challenging. Here, we used data from the National Ecological Observatory Network program to investigate whether ecological relationships among microbial community structure and function and environmental conditions observed at small scales hold at the subcontinental scale. We found that microbial communities were best explained by site-specific conditions. However, when field replicates were averaged, stream physico-chemical characteristics such as pH and temperature emerged as driving factors. This indicates that water quality acted as an environmental filter on microbial communities at the subcontinental scale, but was masked by small-scale patches that created high spatial heterogeneity. Our findings underscore the importance of considering multiple spatial scales to fully understand benthic microbial communities’ role in stream biogeochemistry.

**SCIENTIFIC SIGNIFICANCE STATEMENT:** Microbial communities in streams process materials as water flows toward the ocean. As microbial communities are influenced by environmental conditions, it remains challenging to predict stream microbial metabolism at large spatial scales. In this study, we used the National Ecological Observatory Network (NEON) public database to investigate the drivers of microbial community structure and functions in stream sediments across the USA. This unique dataset revealed high variability in microbial communities among streams that were best explained by local conditions, then by water quality and streambed habitat type. However, stream physico-chemical characteristics emerged as strong predictors of microbial communities when field replicates were averaged and considered as single, large microbial communities. These findings indicate multiple spatial scales must be considered to fully understand benthic microbial communities.

## INTRODUCTION

Streams and rivers are critical components of global elemental cycles, transporting and processing dissolved and particulate matter along the land-to-ocean aquatic continuum (Cole et al. 2007; Maranger et al. 2018; Regnier et al. 2022). Within these ecosystems, metabolism and biogeochemical conditions are driven by microbial communities (Battin et al. 2016). Simultaneously, stream biogeochemical conditions—such as pH, oxygen concentration and organic matter inputs—can also regulate microbial community structure, and there have been substantial research efforts to link microbial communities to specific biogeochemical functions (Bier et al. 2015; Zeglin 2015). Understanding the bidirectional relationship between microbes and their environment is essential for predicting how aquatic ecosystems respond to environmental and climate change. Yet, this remains challenging because of the scale-dependency between microbial communities and environmental variables (Wu and Loucks 1995; LaBrie et al. 2020).

Benthic microbial communities in lotic ecosystems are structured by environmental conditions acting at varying scales. For instance, the immediate overlaying water column controls many physical and chemical conditions such as temperature, organic matter, and nutrient delivery (Battin et al. 2003, 2016; Akinwole et al. 2021). Benthic microbial communities are also influenced by the physical structure of their local habitat, including substrate type and stream depth (Hullar et al. 2006; Cruaud et al. 2020; Gao et al. 2024). These environmental conditions can change at the centimeter to meter-scale, and together, favor high heterogeneity in benthic microbial communities (Wu and Loucks 1995; Clapcott and Barmuta 2010; Briggs et al. 2015). As ecological scale increases, drivers of microbial community structure and function change from being dominated by local stochasticity to larger scale processes such as geological formations and geomorphological and climatological changes (Wu and Loucks 1995; Besemer et al. 2009; Ezzat et al. 2022). Yet, we lack an understanding of the key controls on stream benthic microbial community composition and function at continental scales as most studies have been largely limited to local and regional scales (Zeglin 2015; Hosen et al. 2017; Ruiz-González et al. 2019). In turn, this limits our ability to predict microbial responses to broader climatic changes. A first step in addressing this challenge is to determine whether universal biogeochemical drivers of lotic microbial communities can be identified from existing datasets approaching the continental-scale.

Here, we used data from 23 National Ecological Observatory Network (NEON) streams sampled in 2019 to relate patterns in microbial diversity and function to biogeochemistry across the United States, from Alaska to Puerto Rico. NEON sites can be classified in six different groups based on characteristic water chemistry signatures (Edmonds et al. 2022). As such, we aimed to understand if these water quality groups translated to patterns in microbial communities, as water chemistry has been shown to be an important driver of microbial community structure up to regional scales (Battin et al. 2016; Crowther et al. 2019). Specifically, we sought to answer the following research questions: 1) how do benthic microbial community structure and functional gene diversity vary at the subcontinental scale? 2) is this variation, if present, correlated with patterns in stream biogeochemistry? By using a subcontinental-scale dataset, we can assess the relative contribution of local and regional drivers (e.g., water temperature, biogeochemical conditions, sediment properties) to large-scale patterns in microbial community structure and function.

## METHODS

We leveraged data collected by NEON from 24 wadeable streams (**Table S1**) across the United States, from Puerto Rico to Alaska (**Figure S1**). These streams span 16 of the 20 ecoclimatic domains used by NEON when determining site placement (Hargrove and Hoffman 2004). We targeted sampling efforts between 2019/01/01 and 2019/12/31. This timeframe was selected for even sample coverage across sites; one site (OKSR) was excluded from analysis due to low sequencing yield in 16S rRNA gene and metagenomic data. Land cover varies by site, but the majority of NEON sites are located on smaller streams not highly impacted by anthropogenic activity. Site characteristics such as latitude and longitude, watershed area, and elevation were obtained from the NEON site descriptions (<u>neonscience.org/field-sites/explore-field-sites</u>). Streambed slope was collected from NEON thalweg surveys (DP4.00131.001) and land cover was taken from the NEON Aquatic Watershed shapefile (neon.maps.arcgis.com). Benthic quality-controlled 16S rRNA gene sequences (DP1.20280.001, 2019/01-2019/11), shotgun metagenomic sequences (DP1.20279.001, 2019/06-2019/08), and environmental data (2019/01-2019/11) were downloaded from the NEON data portal using the R package NeonUtilities (Lunch et al. 2024).

### Sampling

The detailed strategy for benthic microbial sampling is available on NEON website (NEON.DOC.003044). Briefly, benthic microbes were sampled three times per year (roughly spring, summer and fall). Only benthic samples taken during the summer sampling event were analyzed for metagenomics. Scrub sampling was performed on rocks (epilithon) and wood (epixylon) to collect the attached biofilm. Biofilms were then resuspended in 125 to 140 mL of 0.2 μm filtered deionized water and then at least 50 mL (or until filter got clogged) were filtered on 0.22 μm filters (Sterivex). Grab samples were collected for sand (epipsammon), silt (epipelon) and plant surfaces (epiphyton) samples. Filters and whirl-pak bags with sample materials were flash-frozen on dry ice in the field and stored at -80 °C until analysis. DNA extraction and sequencing protocols can be found on NEON website (BMI_dnaExtractionSOP, BMI_16Sv4v5_SOP, and BMI_metagenomicsSequencingSOP) and in the SI.

### Bioinformatics

#### 16S rRNA gene

Benthic 16S rRNA gene sequences samples encompass 23 benthic NEON sites (i.e., all but OKSR), totaling 459 samples analyzed in this study. Sequences were analyzed using the dada2 pipeline v1.22.0 (Callahan et al. 2016) in R v4.1.2 (R Core Team 2022). Read quality of each sequence run was visually inspected using quality profiles, and the forward and reverse reads were trimmed to 275 and 250 nucleotides, respectively. Primers were removed using the trimLeft() argument in the filterAndTrim() function. After filtering for quality, the average number of reads was 22,324 per sample. Amplicon Sequence Variant (ASV) tables were produced following the dada2 pipeline including the removal of chimera sequences with an additional step to collapse together sequences that are only different in length (collapseNoMismatch(); minOverlap = 300) to minimize false singletons. Due to the low quality of samples produced by a specific sequence run, we used a threshold of a minimum 2,000 final reads per sample, resulting in the exclusion of 30 samples. Taxonomy assignment was performed at the Genus level with SILVA v138 using dada2’s ‘assignTaxonomy’ function (Wang et al. 2007). Sequences that were unclassified (i.e., no Kingdom assigned; 2567 ASVs) or assigned to eukaryotes or chloroplasts were removed, resulting in 63,813 prokaryotic ASVs. Although not used in this study, surface samples were also curated (see SI) and the final ASV table is publicly available (https://doi.org/10.5281/zenodo.15490674).

#### Metagenomic data

Benthic shotgun metagenomic samples underwent the following bioinformatic pipeline. Paired-end reads were filtered using BBDuk v37.25 (<u>sourceforge.net/projects/bbmap</u>) to remove adapters (k = 12, mink = 11) using normal mode with recommended flags “tpe” and “tbo”. Read pairs for ‘dnaSamples’ corresponding to a single field sample were co-assembled using SPAdes v3.13.0 (Bankevich et al. 2012) without error correction. Filtered reads were aligned to the metagenome assemblies using Bowtie2 v2.4.4 (Langmead et al. 2019) and SAMtools v1.9 (Li et al. 2009) to generate coverage bam files. Contigs greater than 500 basepairs were processed in Anvi’o v7.1 (Eren et al. 2015) for each metagenome assembly. Briefly, anvi-gen-contigs-database was used to generate contig databases and call genes with Prodigal v2.6.3 (Hyatt et al. 2010). Metagenome assemblies were functionally annotated with KOfam, a customized HMM database of KEGG Orthologs (Aramaki et al. 2020), Pfam (Mistry et al. 2021), and NCBI’s database of Clusters of Orthologous Groups of proteins (Tatusov et al. 2003). Next, anvi-profile-blitz was used to export gene-level coverage and detection statistics from the contig databases and coverage bam files. Lastly, functional annotations from all annotation sources were exported from Anvi’o and combined with gene-level statistics into gene-by-sample tables using the custom script process_blitz_neon.R (https://zenodo.org/records/17154747). Gene coverages were normalized based on reads per kilobase per million mapped (RPKM) which corrects differences in both sample sequencing depth and gene length (Konwar et al. 2015). Prodigal gene calls were also annotated using DIAMOND v2.0.15 against NCycDB, a curated database of nitrogen cycling genes (Tu et al. 2019).

#### Environmental data

Environmental data includes grab samples of chemical properties of surface water (DP1.20093.001), continuous discharge (DP4.00130.001), photosynthetically active radiation at the water surface (DP1.20042.001), sediment chemical and physical properties (DP1.20194.001), water temperature (DP1.20053.001), and sensor-based measurements of dissolved oxygen, pH, conductivity, and turbidity (DP1.20288.001). For grab samples, concentrations below the published detection limit were assigned a value of half the detection limit.

Environmental data was aligned with microbial sampling dates by finding the nearest sampling period in time. Some variables were then either combined or removed from the dataset due to significant autocorrelation with other variables (see SI for details). The final dataset contains 35 variables, including 11 watershed and sampling location characteristics, 19 water column variables, and five sediment variables. To assess the influence of water quality using lower dimensionality, we classified sites into one of six water quality groups as determined by Edmonds et al. (2022).

#### Statistical analysis

As sequencing depths and the number of replicates varied among sites for the 16S rRNA gene data, we pooled all field replicates together, computed 1000 different rarefied ASV tables per sample (rrarefy function, vegan, Oksanen et al. 2010) and then averaged them. This enabled us to calculate total richness and Shannon Diversity Index on comparable sites and avoided pseudoreplication when looking at relationships with environmental drivers. Functional diversity metrics were also calculated from gene-by-sample tables (*diversity* function, *vegan*). To ordinate samples based on their compositional dissimilarity, we performed non-metric multidimensional scaling (NMDS) with Bray-Curtis distances (*metaMDS* function, *vegan*) on both metagenomic and 16S rRNA gene data using *phyloseq* v1.50.0 (McMurdie and Holmes 2013). We then projected environmental variables that were significant at p-value < 0.01 on 16S rRNA gene data (*envfit* function, *vegan*). Lastly, statistical differences in taxonomic and functional composition among groups (substrate type, water quality group, NEON site) were tested using PERMANOVA (*adonis2* function, *vegan*) with 999 permutations (Anderson 2001). PERMDISP (*betadisper* and *permutest* functions, *vegan*) was used to test for uneven dispersion of groups (Anderson 2006).

## RESULTS

### Stream benthic communities

Microbial community structure and function composition in benthic streams were best described by local conditions, not water chemistry. Indeed, community structure (**Figure 1A and B**) was best explained by site (PERMANOVA, R^2^ = 0.31, p < 0.001), followed by water quality group (R^2^ = 0.08, p < 0.001) and substrate type (R^2^ = 0.06, p < 0.001). Alpha diversity metrics were generally not well predicted by sites, water quality groups and substrate types, except for the Shannon index which was well constrained by substrate types (**Table S2; Figure S2**). Although there were fewer samples for functional diversity (**Figure 1C and D**), microbial functions were still best explained by site (PERMANOVA, R^2^ = 0.37, p < 0.001), though substrate type (R^2^ = 0.23, p < 0.001) and water quality group explained slightly more variation (R^2^ = 0.14, p < 0.001). Site, substrate type, and water quality group were all significant predictors of the number and diversity of microbial functions (**Table S2; Figure S3**). A general summary of microbial community composition (**Figure S4 to S8**) and metagenomic samples (**Figure S9 and S10**) can be found in the supplementary information.

**Figure 1.**
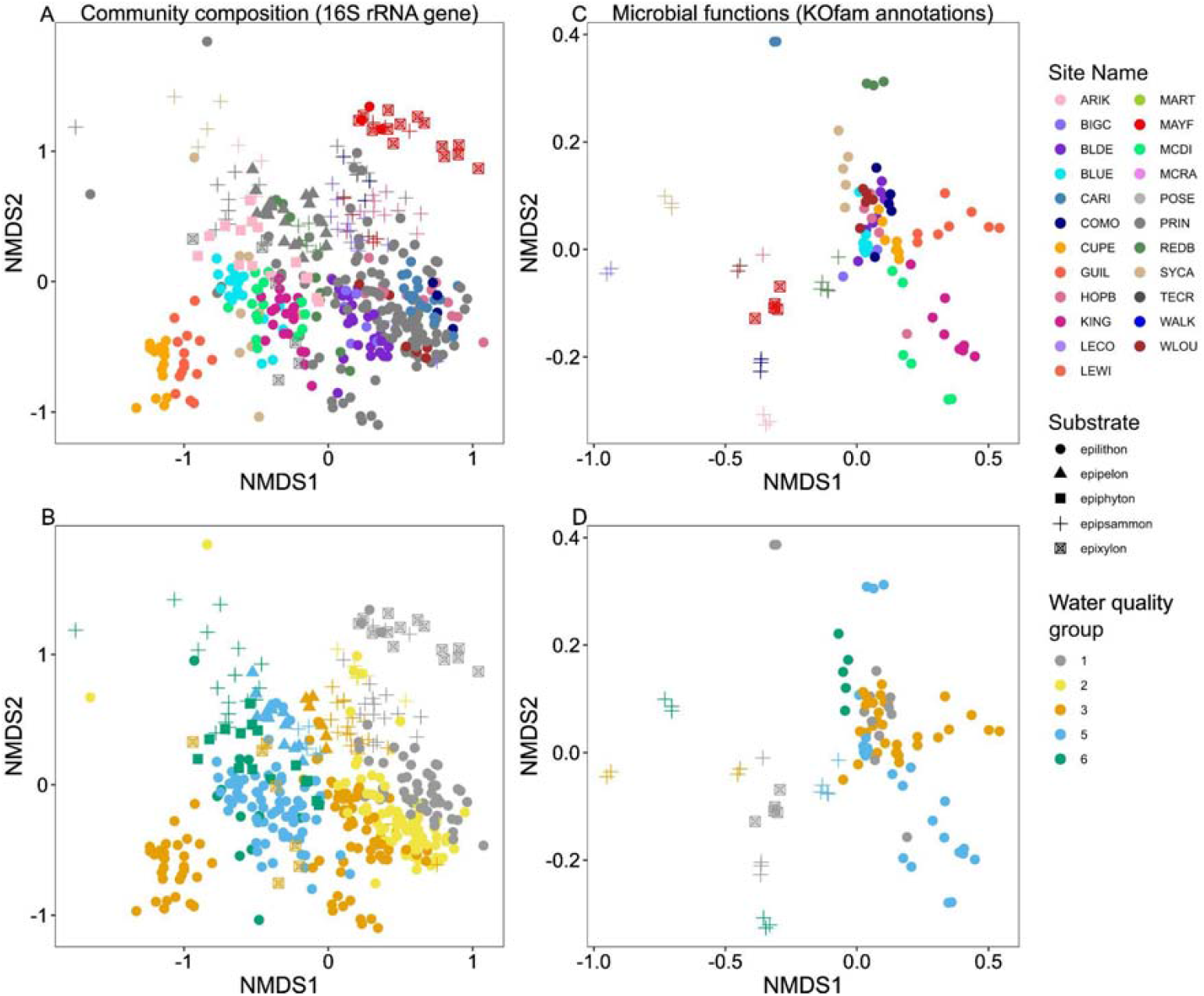
Non-metric multidimensional scaling (NMDS) ordinations showing variation in microbial community composition and function using the 16S rRNA gene (A and B) and KOfam annotations (C and D). Panels A and C are color-coded as a function of site name, B and D, as a function of water quality groups. For all panels, shapes represent substrate types.

### Interrelationships between pH and water quality, and benthic communities

Beyond local conditions, we investigated which environmental variables correlated with community structure across substrate types and within epilithon samples by averaging field replicates. Using NMDS, we displayed samples using either water quality groups (**Figure 2**) or sites (**Figure S11**) and projected environmental variables that were statistically significant. For all substrate types combined, and in epilithon only, the top environmental variables were sediment pH, water quality groups, and water column dissolved inorganic carbon (**Table S3**). Latitude and water temperature were also important for epilithon samples and had opposite effects to each other (**Figure 2B**). Overall, these results show that water quality groups and pH are significantly related to benthic community structure.

**Figure 2.**
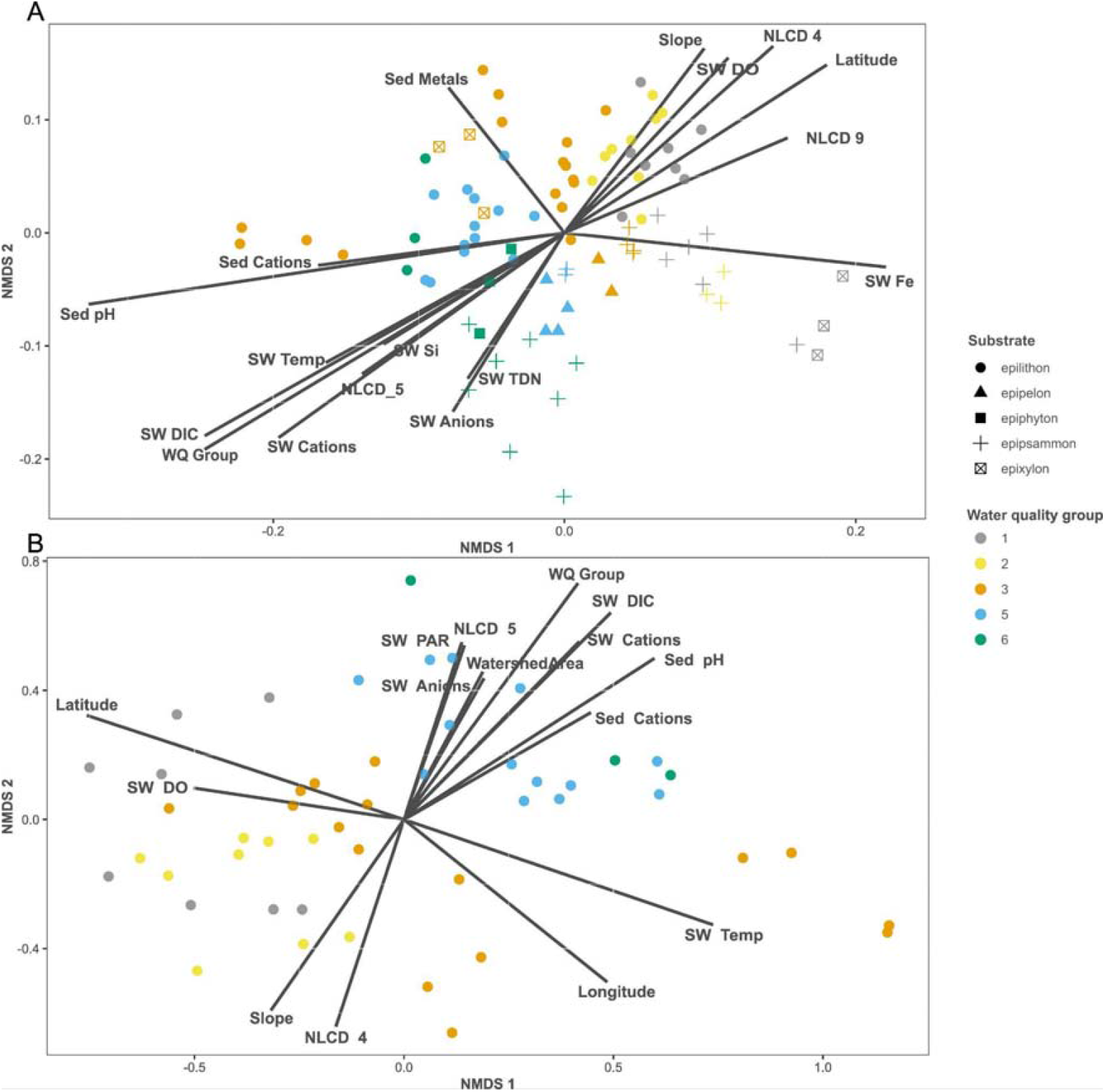
Non-metric multidimensional scaling (NMDS) plot showing variation in microbial community structure (16S rRNA gene) for all substrate types (A) and epilithon only (B) after averaging field replicates. Environmental variables that were statistically significant (p-value < 0.01) were passively projected on the ordination plots using the envfit function. Colors represent water quality groups and shapes, substrate type.

### Abundance of genes related to nitrogen and methane cycling genes vary by substrate type

Using epilithon and epipsammon metagenomic samples, we found that nitrogen- and methane-related genes varied among substrate types (**Figure 3**). Specifically, the average RPKM related to nitrogen was lower in epilithon than in epipsammon, both overall and in site-specific pairwise comparisons (**Figure 3A**), but many nitrogen cycling pathways were ubiquitous across sites and substrates. Anaerobic ammonium oxidation (i.e., anammox), however, was found at only one site (REDB epipsammon) in low abundance (not visible in figure 3A). Across streams, genes associated with heterotrophic processes (e.g., denitrification and dissimilatory nitrate reduction to ammonium [DNRA]) dominated. Taken together, genes associated with nitrogen recycling were more abundant than nitrogen removal. Genes related to methanogenesis were always more abundant than genes related to methane oxidation (**Figure 3B**). While methanogenesis and methane oxidation-related genes covaried in epipsammon samples, they did not in epilithon samples, where methane oxidation-related genes were almost absent.

**Figure 3.**
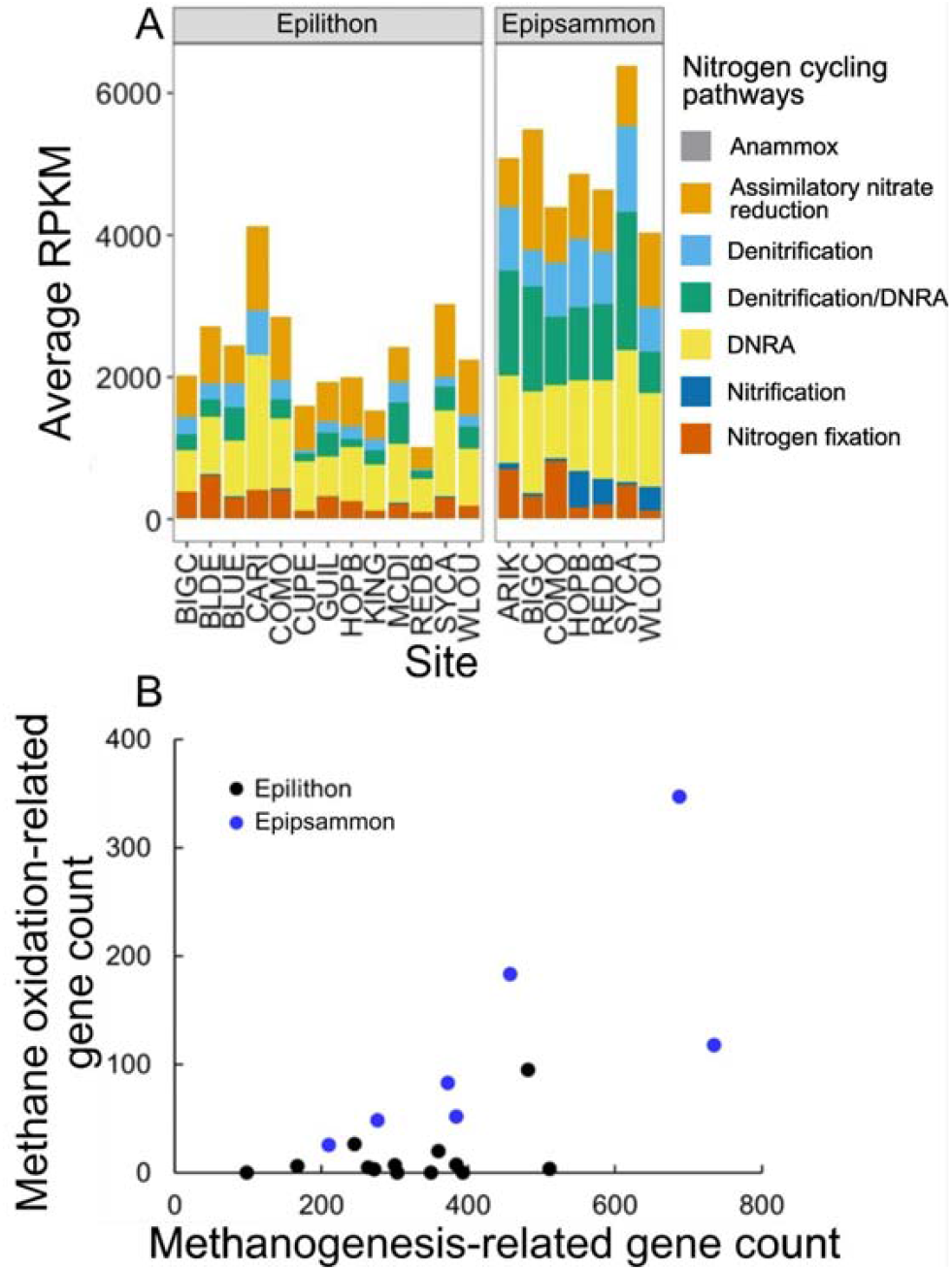
Abundance of nitrogen-related genes as a function of sites and substrate types where colors represent nitrogen cycling pathways (A). Methanogenesis-related and methane oxidation-related genes for epilithon (black) and epipsammon (blue) substrate types (B). In epipsammon, the two methane pathways covaried, but not in epilithon. Anammox: anaerobic ammonium oxidation; DNRA: dissimilatory nitrate reduction to ammonium; RPKM: reads per kilobase per million mapped

## DISCUSSION

Using publicly available NEON microbial sequencing data and biogeochemistry data, we present the first subcontinental patterns in microbial taxonomic and functional diversity in streams located across the USA. When considering field replicates individually, we found that water quality was not the main driver of community structure. However, when field replicates were averaged, water quality groups as an integrative variable became a strong predictor of community structure and function. More specifically, pH and average temperature were strong predictors of community structure as observed in soil and other stream studies (Sutton and Findlay 2003; Lauber et al. 2009; NiñoLGarcía et al. 2016; Philippot et al. 2024). Previous studies observing ecologically significant relationships between geochemical parameters and benthic microbial communities have generally been limited to highly impacted streams (Zhou et al. 2021) or extreme environments (Ezzat et al. 2022). Thus, although the water quality groups used in this study are coarse, our findings suggest that broad-scale patterns in water quality act as environmental filters for benthic microbial communities at subcontinental scales even in relatively pristine systems, with implications for understanding stream microbial processes at the continental scale.

### Substrate type influences genes related to biogeochemical transformations

Across all sites, community structure and function were best explained by location, but could also be explained by substrate type. This supports the idea that habitat structure influences microbial communities in aquatic ecosystems, as observed at smaller scales (Besemer et al. 2009; Singer et al. 2010; Cruaud et al. 2020; Sun et al. 2023). For instance, streambed composition controls the life-cycle of biofilms (Chen et al. 2022) and influences oxygen exchange between running water and hyporheic flow (Jones and Holmes 1996; Mueller et al. 2013). Oxygen availability likely contributed to the contrasting microbial communities, and specifically genes related to nitrogen metabolic pathways, between epipsammon and epilithon substrates. Observed differences in nitrogen cycling microbial communities were likely related to redox gradients and although anoxia can develop in epilithic biofilms (Anderson-Glenna et al. 2008; Flemming et al. 2016), it appeared to be less common in our dataset, potentially as a result of the sampling strategy—grab samples were used for epipsammon in which oxic-anoxic gradients can be present at the centimeter scale (Briggs et al. 2015). In epipsammon samples, nitrification genes coexisted with denitrification and DNRA, as did methanogenesis and methane oxidation-related genes. As these metabolic pathways are dependent on oxic-anoxic coexistence, it suggests that spatial complexity (i.e., patches) of oxygen availability had a stronger influence on microbial communities than nutrient availability. Patches of oxygen availability were likely more abundant and in closer proximity in epipsammon due to finer substrates than in epilithon samples.

### Patches across spatial scales

Averaging community structure on a site–substrate type basis revealed the strong influence of water quality groups, pH and temperature on microbial communities. This implies high spatial heterogeneity in which small-scale patches within stream reaches obscured the influence of biogeochemical conditions (Akinwole et al. 2021; Bier et al. 2023). Indeed, the choice of scale strongly influences our ability to observe patterns (Wu and Loucks 1995) where small-scale processes may appear as stochastic variation at larger scales, and likewise, large-scale relationships may dissipate at smaller scales. Patches, however, are not limited to small-scale features; they can also appear at larger scales, creating nested processes (Wu and Loucks 1995). For instance, Puerto Rican CUPE and GUIL sites did not differ much from other sites in terms of water and sediments biogeochemical conditions (**Figure S12**), but had unique epilithon communities, displayed the highest level of unique taxa and among the highest number of metabolic functions. This suggests that local conditions specific to these sites were not well-captured despite the broad NEON scope. Local enrichments to benthic microbial communities likely created patches of high diversity in both CUPE and GUIL sites—Puerto Rico may thus represent a large-scale patch up to ∼10,000 km^2^ that is both nesting smaller scale patches and is nested within the larger system (**Figure 4**). Revealing the ecological importance of patches at the subcontinental scale was possible given the high spatial sampling effort—both within reach and across ecoregions—of the NEON program.

**Figure 4.**
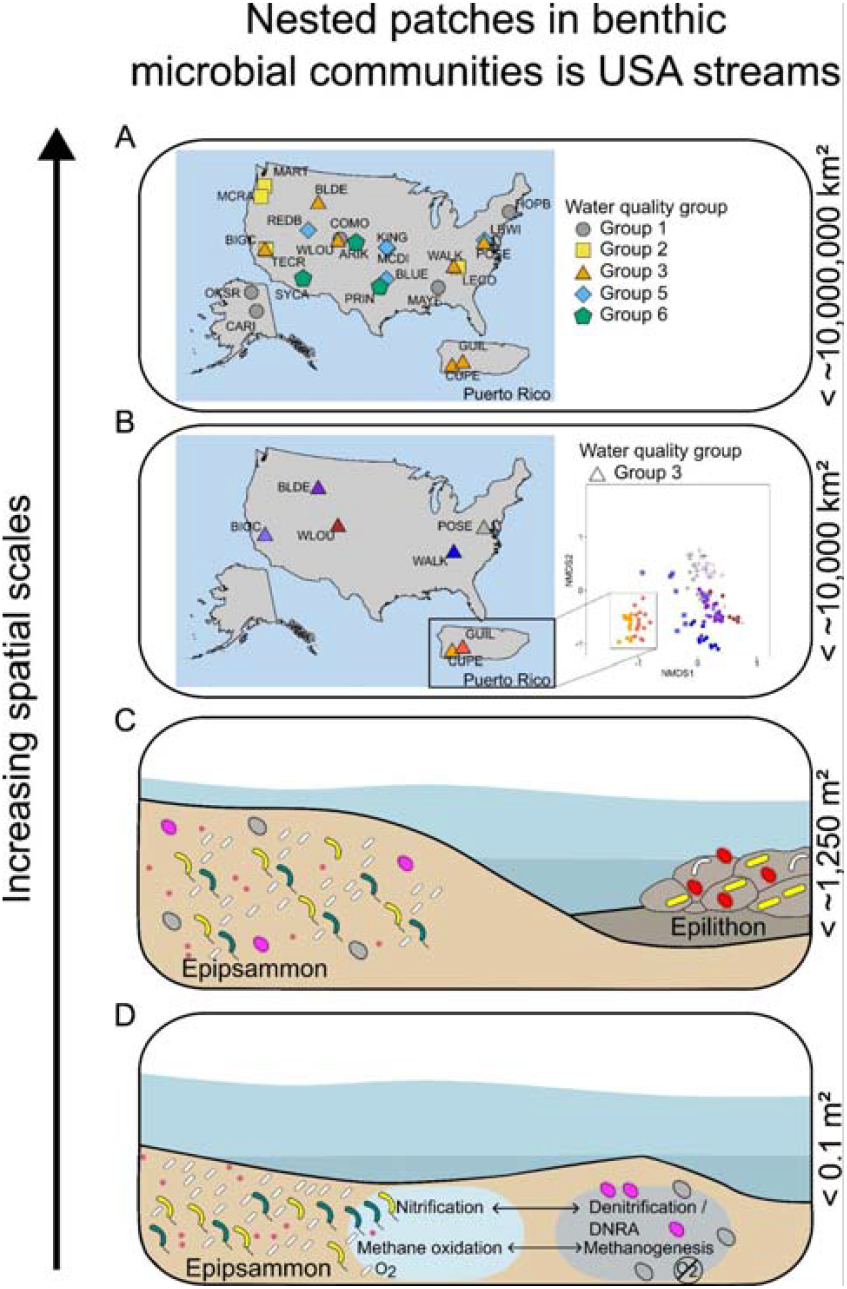
Representation of nested patches across spatial scales, including at the subcontinental scale with water quality groups (A, < ∼10,000,000 km^2^), Puerto Rico CUPE and GUIL sites (B, < ∼10,000 km^2^), among substrate type (C, < ∼1,250 m^2^), and within a grab sample (D, < 0.1 m^2^). Symbols in A and B represent water quality groups as defined by Edmonds et al. (2022), and represent microbial communities in C and D. The inset in B uses the same NMDS as figure 1A but only displays sites for water quality group 3; colors are shared between the map and the NMDS.

## CONCLUSION

Open science provides a segue from spatially-limited field observation programs to large-scale studies. By expanding spatial extents to subcontinental scales, environmental gradients and ecological niches are more diverse, allowing robust assessments of our fundamental understanding of aquatic ecosystems and ecological theories. While biogeochemical conditions can be the main driver of community structure in streams (Peipoch et al. 2019), our subcontinental approach highlighted the greater influence of patches, from small (within reach) to large (Puerto Rico and larger) scales. Although within-reach patches did not relate broadly to water and sediment biogeochemical conditions, presumably because of the scale-dependency of environmental variables (LaBrie et al. 2020), patches are critical for stream ecosystem processes by maximizing microbial biomass and functions through niche partitioning (Cardinale 2011; Battin et al. 2016), ensuring high functional redundancy and resilience (Comte et al. 2013; Pelletier et al. 2020). Our findings suggest that multiple spatial scales—from within a stream reach to continental scale—must be considered to fully understand stream benthic microbial communities and how they impact biogeochemical cycles.

## ACKNOWLEDGEMENTS

This material is based in part upon work supported by the National Ecological Observatory Network (NEON), a program sponsored by the U.S. National Science Foundation (NSF) and operated under cooperative agreement by Battelle. We thank the Ecological Dissertations in the Aquatic Sciences (Eco-DAS)-XIV organized by Paul Kemp and Krissy Remple. We thank Andrew H. Morris for discussions and advice on analyzing metagenomic data. We thank the National Science Foundation (NSF, award OCE-1925796) and Association for the Sciences of Limnology and Oceanography (ASLO) for funding and fostering this collaboration. Part of this work was funded by the Wares and Trottier Space Institute postdoctoral fellowships to RL; the AMTD Waterloo Global Talent postdoctoral fellowship to PCJR and the Friday Harbor Laboratories William Calvin Postdoctoral Fellowship to RLM.

## Data Availability Statement

Data are available in the Zenodo repository at https://doi.org/10.5281/zenodo.15490674

