## Supplementary information for "Small- and large-scale patches shape benthic microbial community structure and function in streams at the subcontinental scale"

### METHODS

*Sampling*

The detailed strategy for surface water sampling is available on NEON website (NEON.DOC.001152). Sampling was performed approximately monthly for surface water microbes and 0.5–2 L was filtered on 0.22 µm filters (Sterivex, Millipore) in the field.

*Nucleic acid extraction and sequencing*

One to eight DNA extractions occurred on a single field sample (Site-Date-Substrate type) provided to the analytical lab, and each of these 'dnaSamples' were sequenced. DNA was extracted using DNeasy PowerBiofilm Kit (Qiagen) for swab samples (epiphyton, epipsammon and epipelon) and DNeasy PowerWater Sterivex™ Kit (Qiagen) for sterivexes (surface water, epilithon and epixylon). After extraction, DNA was quantified using a Quantus Fluorometer with a QuantiFluor ONE dsDNA Kit and quality reviewed following batch- and sample-level quality control criteria. PCR was conducted with primers 515F (5'-GTGYCAGCMGCCGCGGTAA-3') and 926R (5'-CCGYCAATTYMTTTRAGTTT-3') for the amplification of the V4V5 region of the 16S rRNA gene (Parada et al. 2016). High-throughput paired-end sequencing of the 16S rRNA v4v5 region was performed on the Illumina MiSeq platform. Samples were prepared for metagenomic shotgun sequencing using the KAPA Hyper Plus kit (Kapa Biosystems). Final libraries were quantified by qPCR, normalized, pooled into sets of 40 samples, and sequenced on the Illumina NextSeq 550.

*Bioinformatics: 16S rRNA gene*

Surface water (01/2019–12/2019) quality-controlled 16S rRNA gene sequences collected in streams were downloaded from the NEON data portal (DP1.20282.001) on 2022-02-22 using the R package neonUtilities. These samples encompass 23 benthic NEON sites (i.e., all but OKSR). In total, 180 samples were analyzed.

Surface water sequences were analyzed using the dada2 pipeline v1.22.0 (Callahan et al. 2016) in R v4.1.2 (The R Core Team, 2021). Read quality of each sequence run was visually inspected using quality profiles, and the forward and reverse reads were trimmed to 275 and 250 nucleotides, respectively. Primers were removed using the trimLeft() argument in the filterAndTrim() function. After filtering for quality, the average number of reads per sample was 65,873. Amplicon Sequence Variant (ASV) tables were produced following the dada2 pipeline including the removal of chimera sequences and an additional step to collapse together sequences that are only different in length (‘collapseNoMismatch()’; minOverlap = 300), which minimizes false singletons. Due to the low quality of samples produced by a specific sequence run of benthic samples, we used a threshold of a minimum 2,000 final reads per sample, resulting in the exclusion of 10 surface water. Taxonomy assignment was performed at the Genus level with SILVA reference database v138 using dada2's ‘assignTaxonomy’ function (Wang et al. 2007). Sequences that were unclassified (i.e., no Kingdom assigned; 550 samples) or assigned to eukaryotes or chloroplast were removed, resulting in 35,633 ASVs. The final ASV table is publicly available (https://zenodo.org/records/15490674).

*Environmental data*

For surface water and sensor-based data, 93% of sampling dates were less than two weeks different between microbial and environmental samples, and 87% of sediment data was within six months of the microbial sample. Sensor data represents the average of all measurements over the seven days before microbial sampling. Heavy metals and toxic elements in sediment data were combined into a single variable; $Na^{+}$, $Ca^{2+}$, $Mg^{+}$, and $K^{+}$ were combined as cations (sediments and surface water) and $Br^{-}$, $Cl^{-}$, $F^{-}$, $SO_4^{2-}$ were combined as anions (surface water only). One example of environmental variables removed from consideration were components of the carbonate system (i.e., OH-, HCO3-, CO3, alkalinity) in the surface and sediment samples while keeping pH and DIC. Examples of highly correlated variables include using total dissolved nitrogen in place of nitrate and total nitrogen in surface water samples, and using the percent clay in the sediment in place of percent silt, percent sand, and total sediment solids.

### RESULTS

#### Metagenomic data

*Contig assembly and functional gene annotation*

In total, 99 metagenomic sequencing samples were co-assembled by site and substrate type into 22 metagenome assemblies (**Figure S9**). All NEON shotgun sequenced samples used to assemble contigs yielded a total of 812,131,128 read pairs at an average of 8,203,345 read pairs per sample (± 533,052; standard error). Overall, benthic metagenome samples exhibited a wide range of sequencing depths which is affected by DNA amount in the original sample, lab DNA extraction efficiency, and sequencer error (**Figure S9A**). After filtering raw reads, an average of 90.8 ± 0.2% of read pairs remained **(Figure S9A**). After metagenomic assembly, an average of 24.2 ± 1.1% of read pairs within a library were recruited to assembled contigs (5,074,582 contigs total; average contigs/library = 230,663 ± 34,080; **Figure S9A**). The variation in sequencing depth generally corresponded to high variation in assembly length and number of gene calls (**Figure S9B**). Generally, samples with deeper sequencing depths produced assemblies with greater contig N50 values (**Figure S9C**). The percentage of genes that were annotated depended both on the annotation source and the sample substrate type with epilithon samples having higher percent annotation compared to epipsammon or epixylon perhaps due to higher quality assemblies (**Figure S9C&D**).

### Supplementary Figures and Tables

Table S1. Site descriptions for NEON wadeable streams.

| Site Abbreviation | Site Name | Latitude | Longitude | Dominant LULC | Watershed area (km2) | Slope (unitless) |
|---|---|---|---|---|---|---|
| ARIK | Arikaree River | 39.758206 | -102.44715 | Cultivated crops | 2600 | 0.003 |
| BIGC | Upper Big Creek | 37.059719 | -119.25755 | Forest | 11 | 0.005 |
| BLDE | Blacktail Deer Creek | 44.95011 | -110.58715 | Grassland/herbaceous | 38 | 0.018 |
| BLUE | Blue River | 34.444218 | -96.624201 | Grassland/herbaceous | 320 | 0.002 |
| CARI | Caribou Creek | 65.153224 | -147.50397 | Forest | 31 | 0.011 |
| COMO | Como Creek | 40.034962 | -105.54416 | Forest | 3.6 | 0.087 |
| CUPE | Rio Cupeyes | 18.11352 | -66.98676 | Forest | 4.3 | 0.031 |
| GUIL | Rio Yahuecas | 18.17406 | -66.79868 | Forest | 9.6 | 0.016 |
| HOPB | Lower Hop Brook | 42.471941 | -72.329526 | Forest | 12 | 0.026 |
| KING | Kings Creek | 39.105061 | -96.603829 | Grassland/herbaceous | 13 | 0.0001 |
| LECO | LeConte Creek | 35.690428 | -83.50379 | Forest | 9.1 | 0.056 |
| LEWI | Lewis Run | 39.095637 | -77.983216 | Pasture | 12 | 0.008 |
| MART | Martha Creek | 45.790835 | -121.93379 | Forest | 6.3 | 0.025 |
| MAYF | Mayfield Creek | 32.960365 | -87.407688 | Forest | 14 | 0.003 |
| MCDI | McDiffett Creek | 38.945861 | -96.443022 | Grassland/herbaceous | 23 | 0.007 |
| MCRA | McRae Creek | 44.259598 | -122.16555 | Forest | 3.9 | 0.068 |
| OKSR | Oksrukuyik Creek | 68.669753 | -149.14302 | Shrublands | 58 | 0.007 |
| POSE | Posey Creek | 38.89431 | -78.147258 | Forest | 2 | 0.03 |
| PRIN | Pringle Creek | 33.378517 | -97.782312 | Grassland/herbaceous | 49 | 0.002 |
| REDB | Red Butte Creek | 40.783934 | -111.79789 | Forest | 17 | 0.045 |
| SYCA | Sycamore Creek | 33.750993 | -111.50809 | Shrublands | 280 | 0.01 |
| TECR | Teakettle Creek - Watershed 2 | 36.955931 | -119.02736 | Forest | 3 | 0.072 |
| WALK | Walker Branch | 35.957378 | -84.279251 | Forest | 1.1 | 0.033 |
| WLOU | West St Louis Creek | 39.891366 | -105.9154 | Forest | 4.9 | 0.046 |

Note. OKSR site was not included in the analysis but appears in the table for reference.

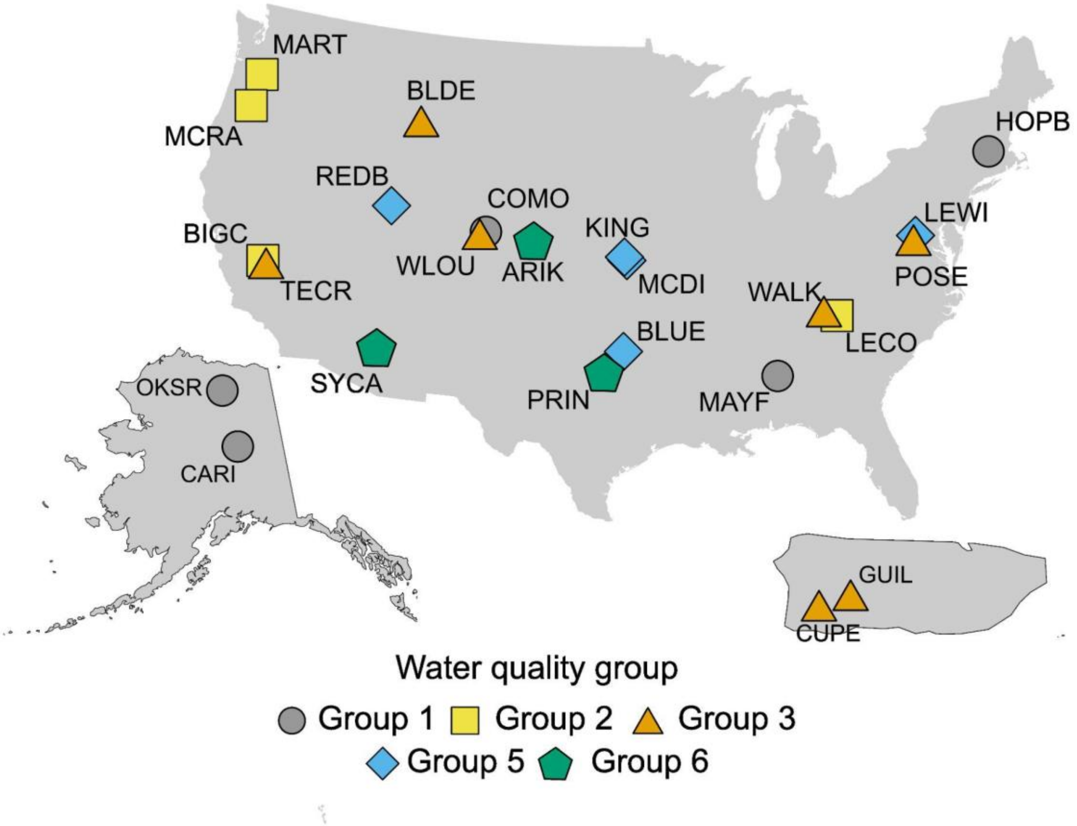


Figure S1. Location of our study sites across the conterminous USA (top), Alaska (bottom left) and Puerto Rico (bottom right). OKSR site was not included in the analysis but appears on the map for reference. Colored symbols refer to water quality groups as defined by Edmonds et al. (2022).

Table S2. Differences in richness and Shannon Diversity Index between sites, water quality groups, and substrate types for community taxonomic composition (16S rRNA gene) and microbial functions (KOfam Annotations) using the Kruskal-Wallis chi-squared test.

| | Richness | | | Shannon diversity index | | |
|---|---|---|---|---|---|---|
| *Composition* | $\chi^2$ | df | p-value | $\chi^2$ | df | p-value |
| Site | 21.849 | 22 | 0.44 | 19.004 | 22 | 0.65 |
| Water quality group | 5.5851 | 4 | 0.23 | 1.2678 | 4 | 0.87 |
| Substrate type | 3.2319 | 4 | 0.52 | 28.315 | 4 | <0.001 |
| *Function* | | | | | | |
| Site | 61.07 | 18 | < 0.001 | 38.39 | 18 | <0.01 |
| Water quality group | 27.19 | 5 | <0.001 | 15.21 | 5 | <0.01 |
| Substrate type | 69.82 | 4 | <0.001 | 71.54 | 4 | <0.001 |

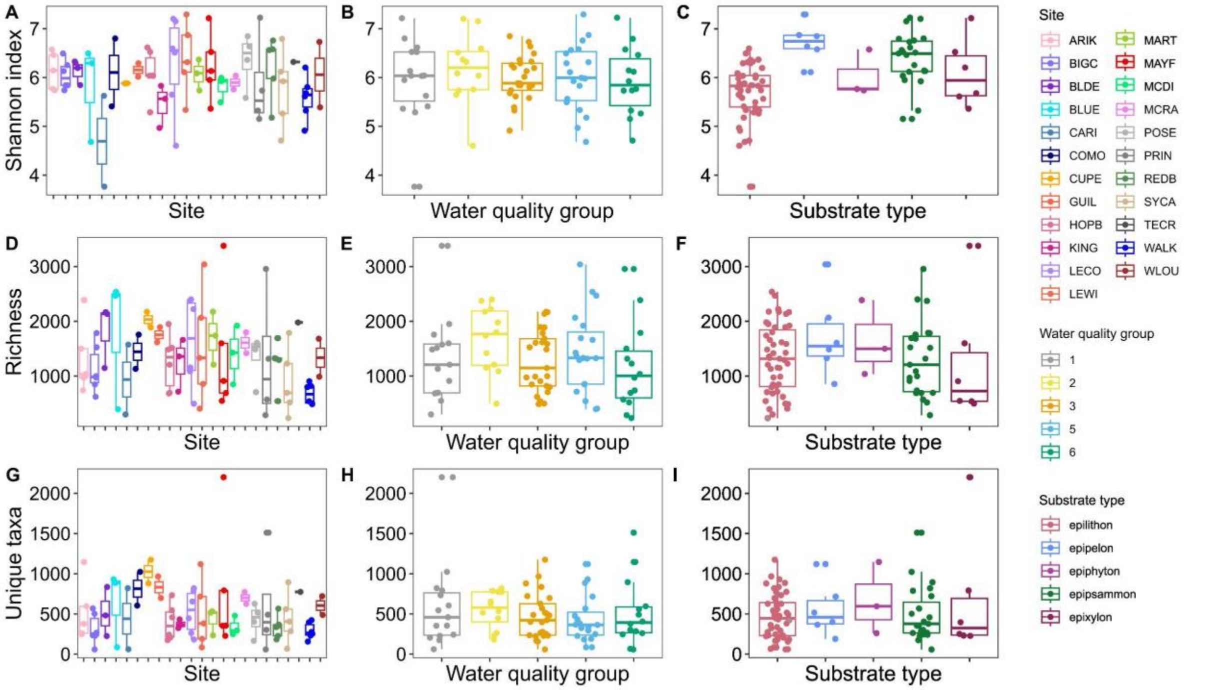


Figure S2. Box and whiskers plots showing variability in Shannon index (top row) and richness (middle row) and unique taxa (bottom row) using 16S rRNA gene sequencing. Samples are colored as a function of site ID (A, D and G), water quality group (B, E and H) and substrate type (C, F and I).

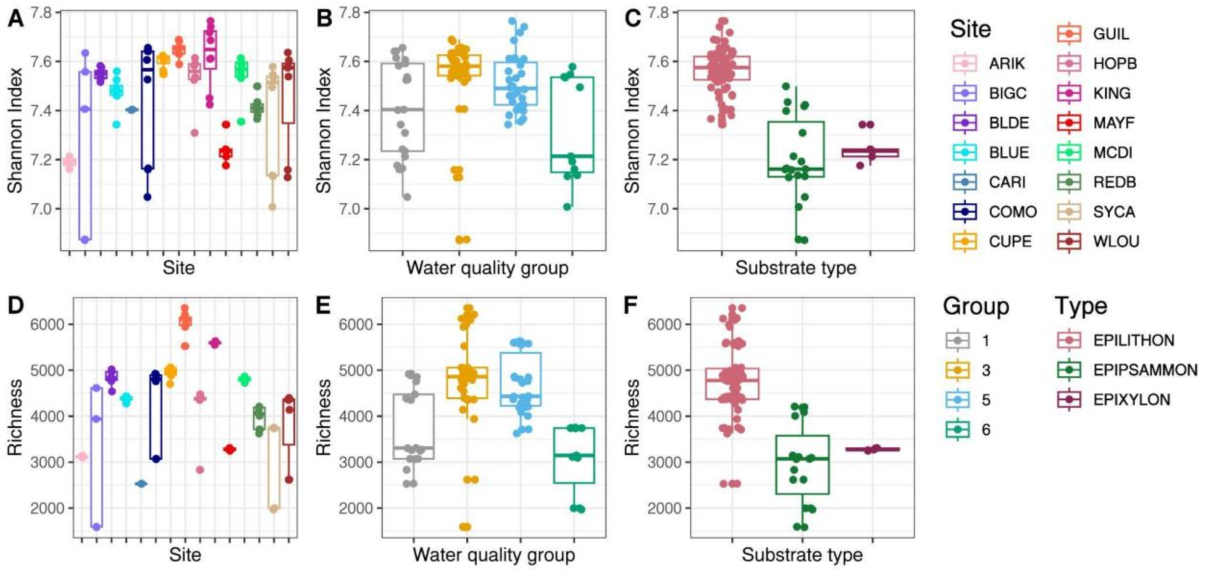


Figure S3. Box and whiskers plots showing variability in Shannon index (top row) and richness (middle row) of annotated KOfam functions from metagenome assemblies. Samples are colored as a function of site ID (A, D), water quality group (B, E) and substrate type (C, F).

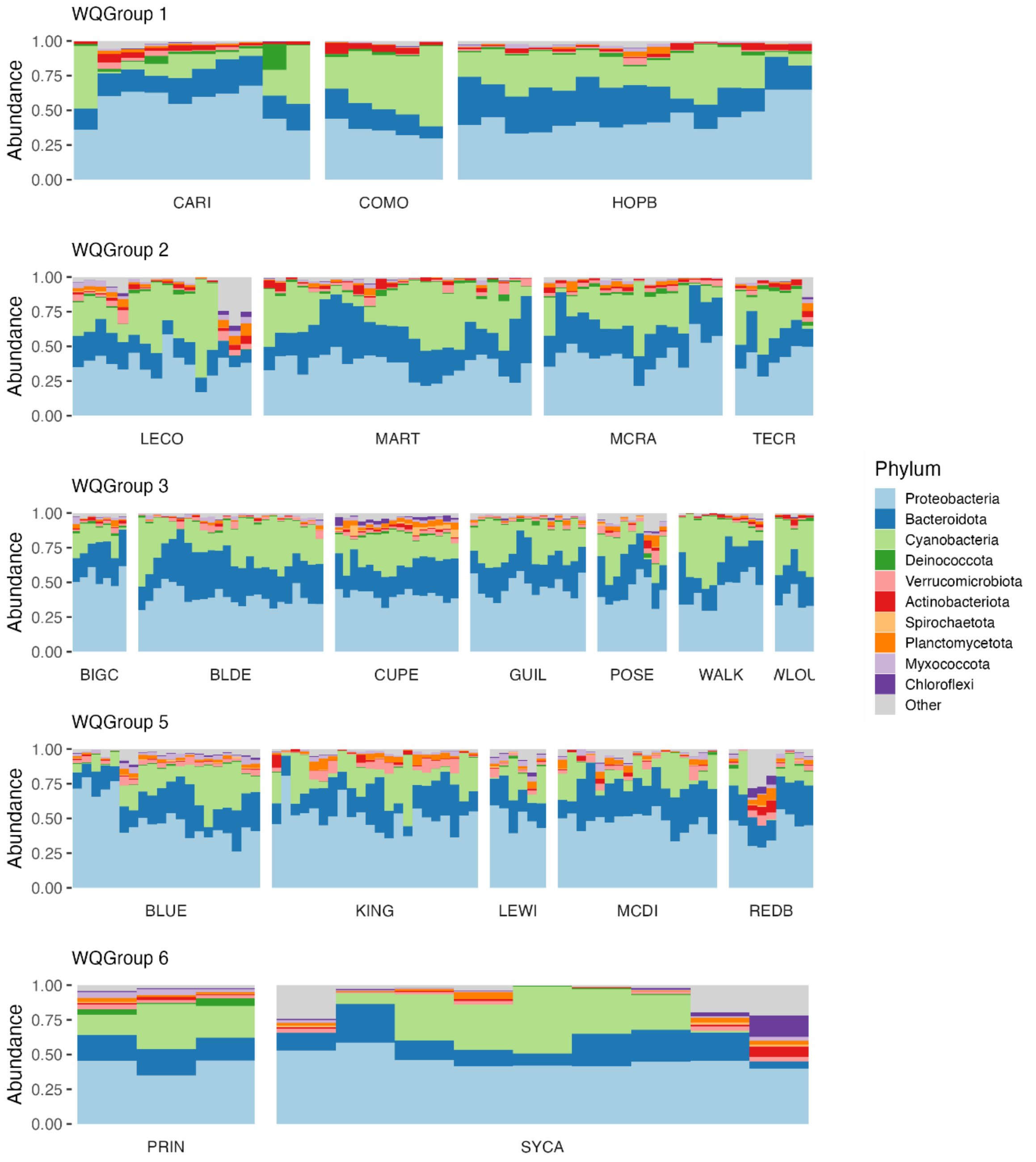


Figure S4. Microbial community composition across epilithon (rocks) sites at the phylum level.

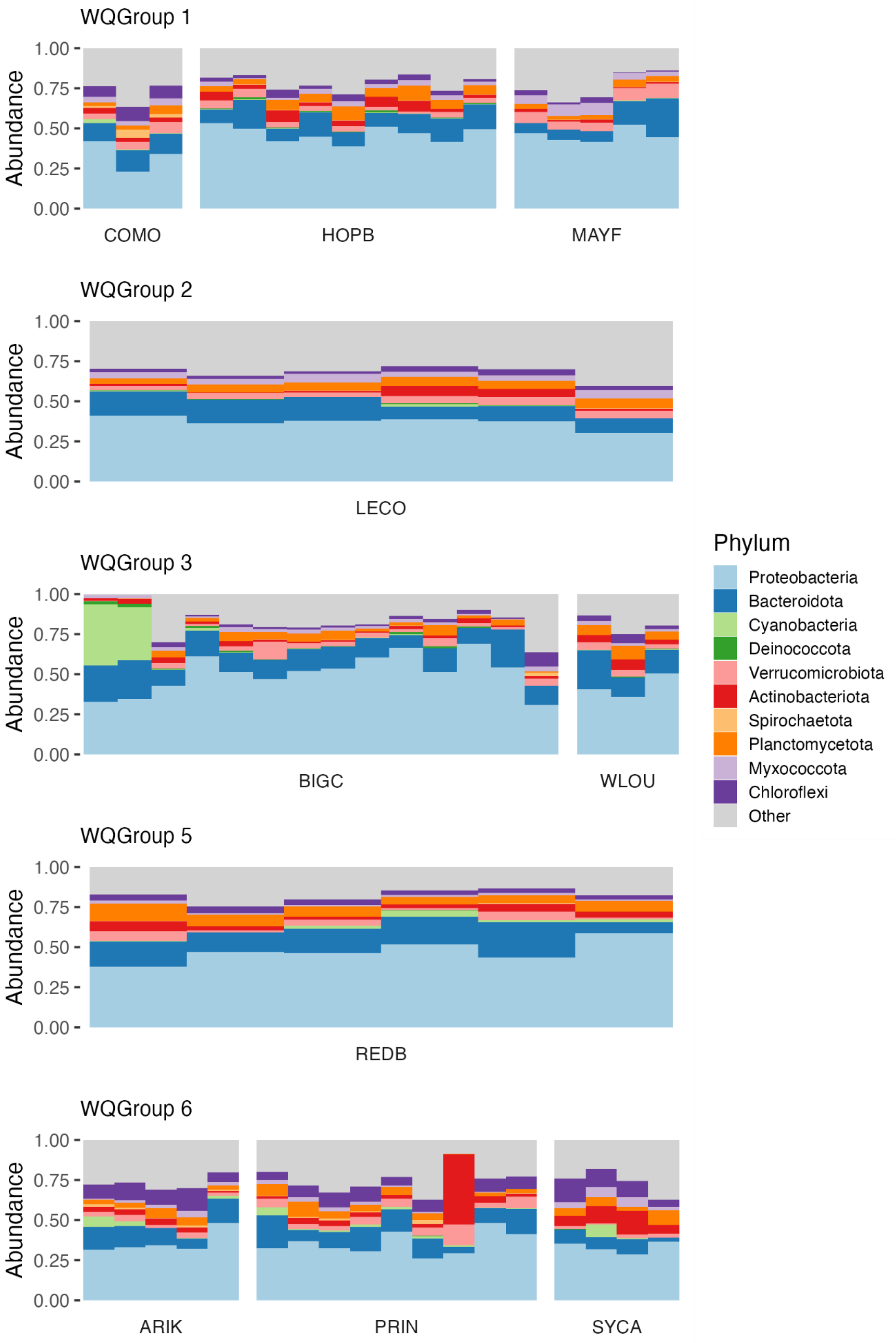


Figure S5. Microbial community composition across epipsammon (sand) sites at the phylum level.

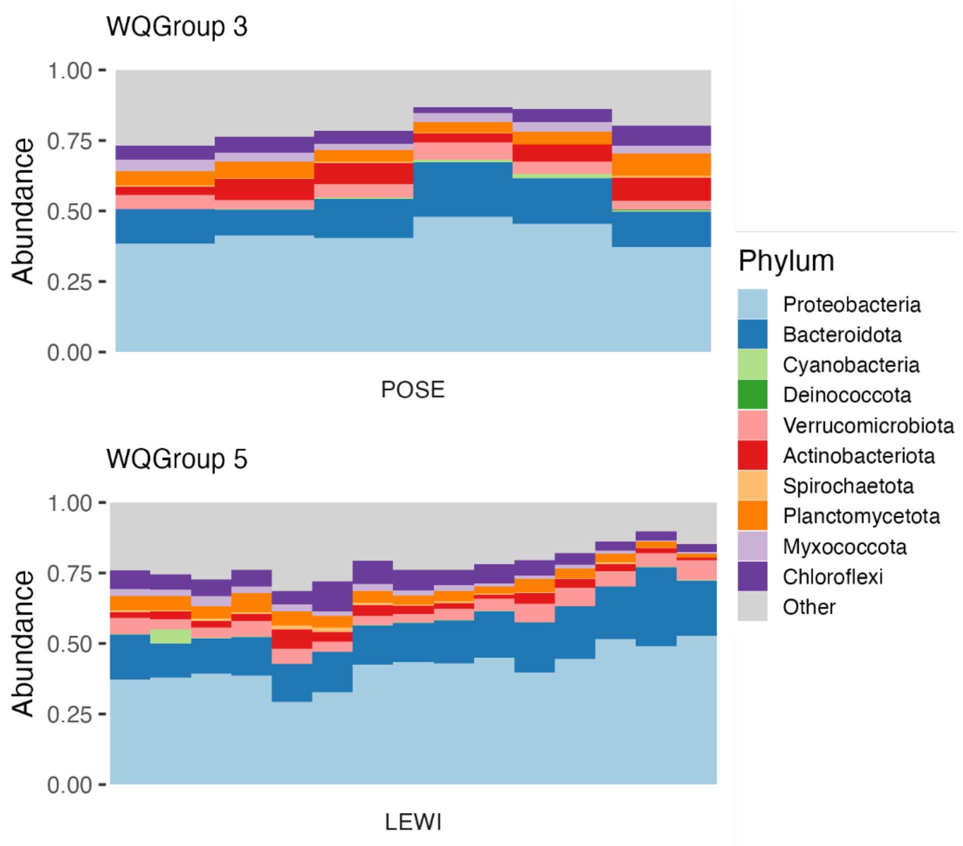


Figure S6. Microbial community composition across epipelon (silt) sites at the phylum level.

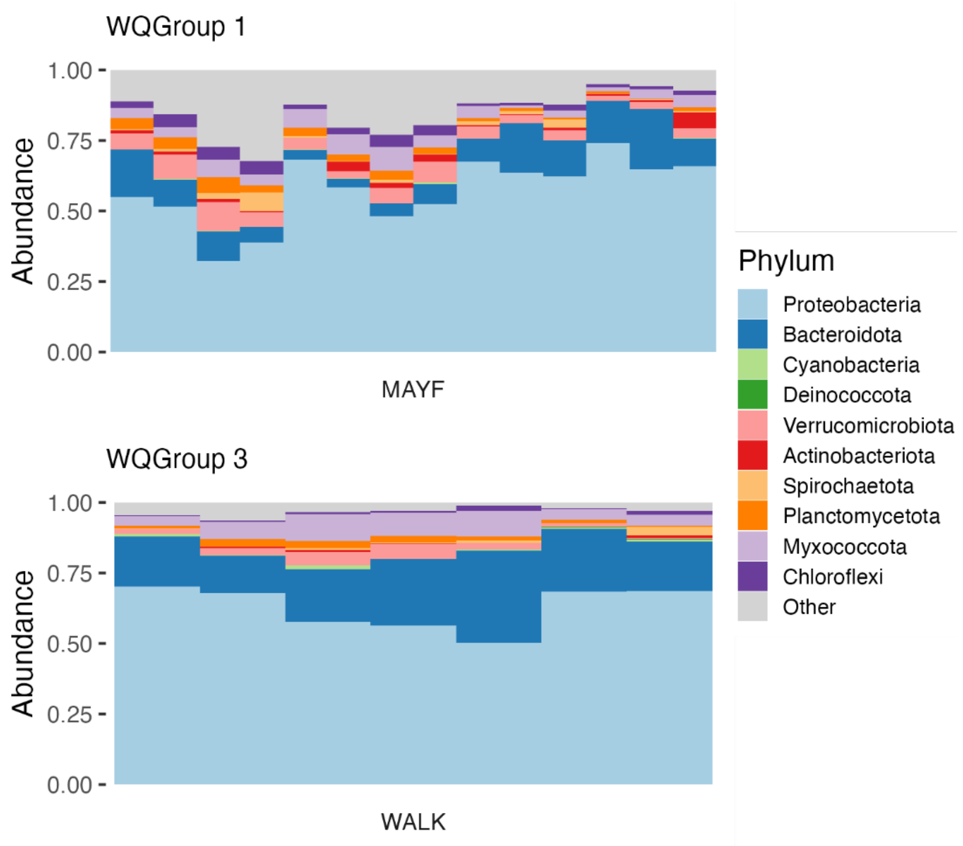


Figure S7. Microbial community composition across epixylon (wood) sites at the phylum level.

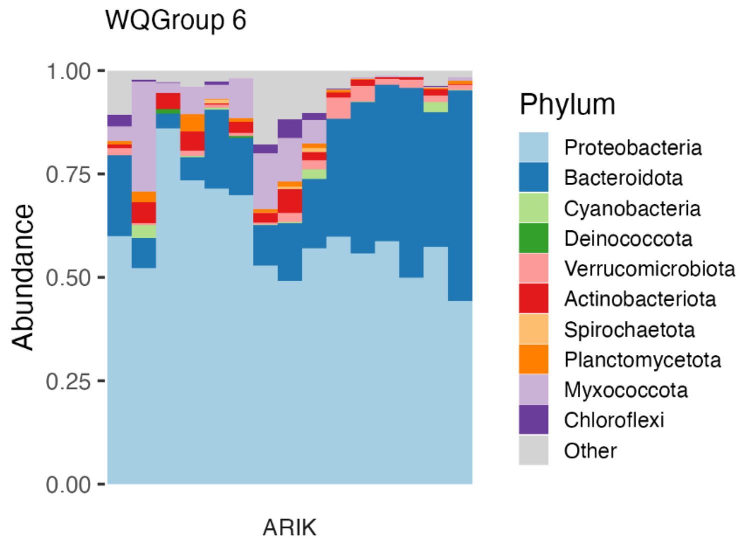


Figure S8. Microbial community composition across epiphyton (plant) sites at the phylum level.

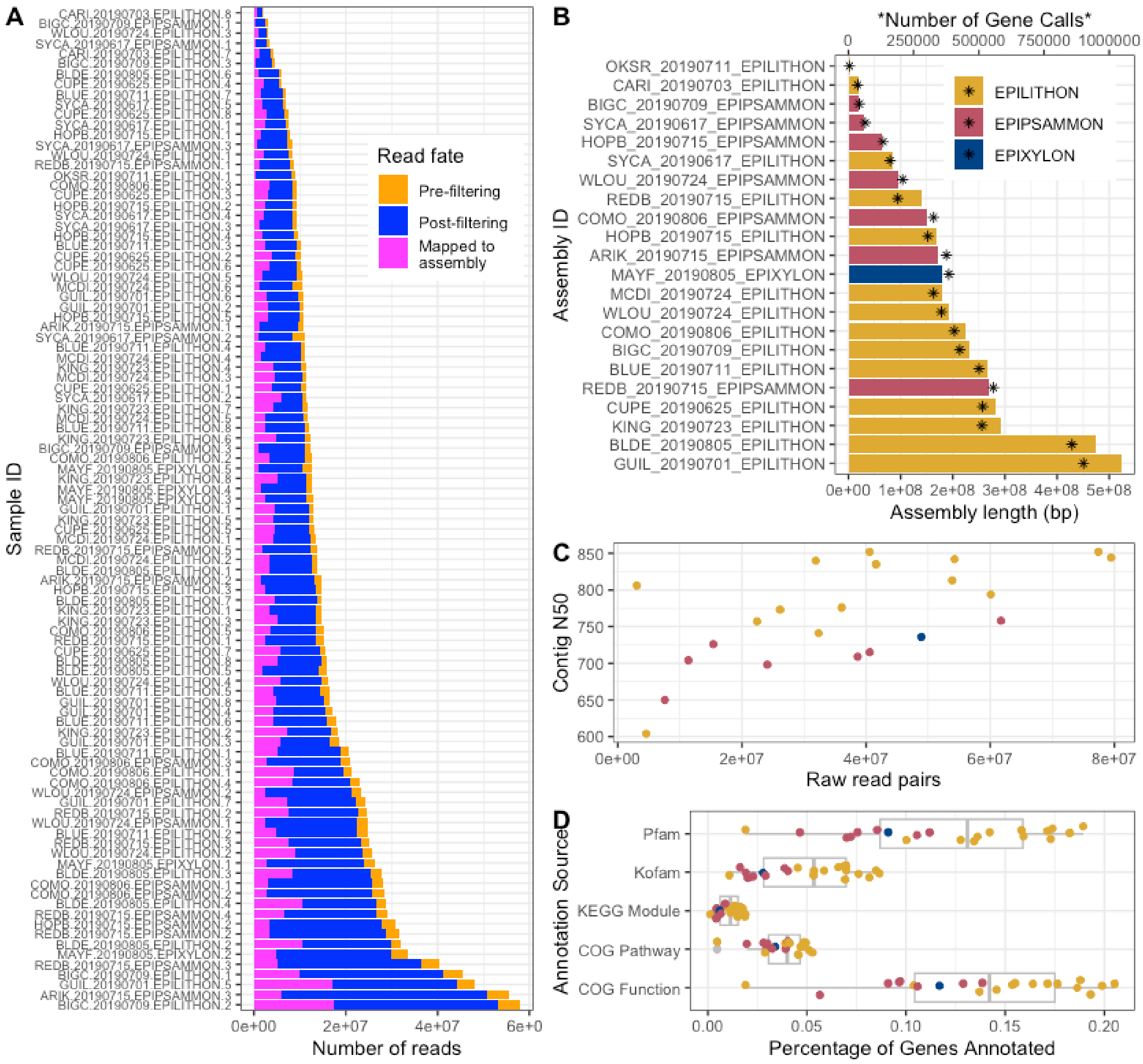


Figure S9. Overview of metagenome reads, assembly, and annotation statistics. A) Number of reads in each NEON metagenome sample that remained after filtering and that successfully mapped to an assembly (listed in B). B) Number of gene calls and length of assemblies of co-assembled NEON metagenome samples. Replicate samples were co-assembled by site and substrate type. C) Relationship between contig N50 and initial raw reads per metagenome assembly. Points are colored by substrate type. D) Percentage of genes annotated in each assembly by annotation source.

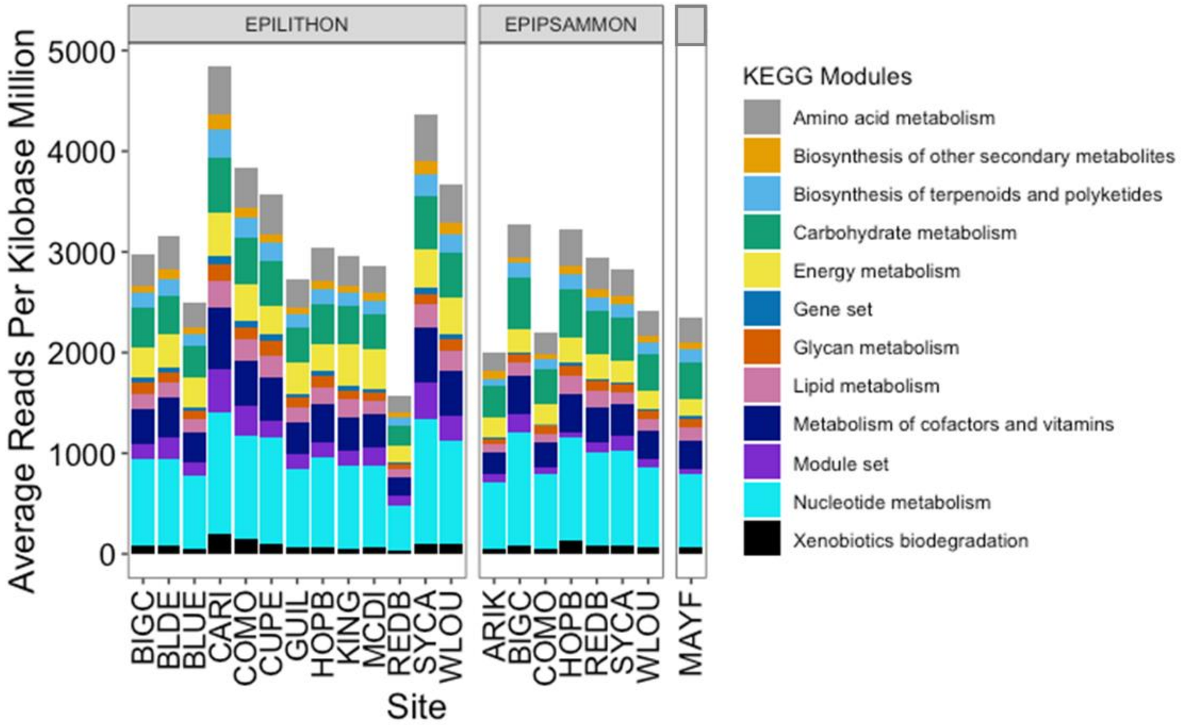


Figure S10. Functional annotation of the metagenome presented as a function of sites and substrate types. The site MAYF is the epixylon (wood) sample and is presented here for reference.

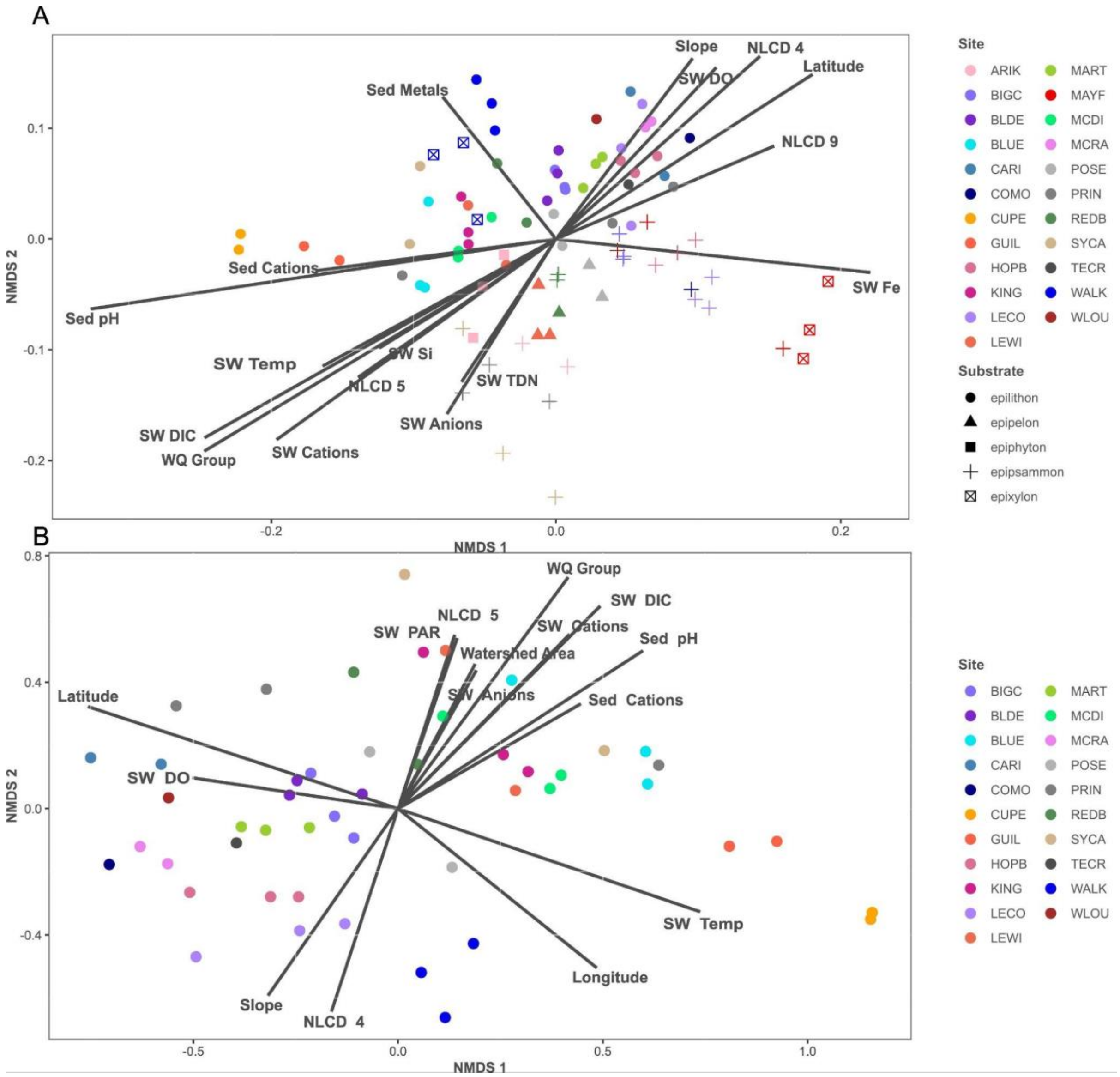


Figure S11. Non-metric multidimensional scaling (NMDS) plot showing variation in microbial community structure for all substrate types (A) and epilithon-only (B). Environmental variables that were statistically significant (p-value<0.01) were passively projected on the ordination plots using the envfit function. Colors represent site ID and shapes, substrate type.

| Substrate | Variable | $R^2$ |
|---|---|---|
| All substrate type | Sediment pH | 0.70 |
| | Water quality groups | 0.62 |
| | Water column DIC | 0.59 |
| | Water column Cations | 0.45 |
| | Latitude | 0.34 |
| | Water column Iron | 0.31 |
| | NLCD 4 | 0.30 |
| | Water column Temperature | 0.25 |
| | Slope | 0.23 |
| | Water column Dissolved Oxygen | 0.23 |
| | NLCD 5 | 0.22 |
| | NLCD 9 | 0.19 |
| | Water column Anions | 0.19 |
| | Sediment Cations | 0.19 |
| | Water column Silicon | 0.16 |
| | Sediment Metals | 0.14 |
| | Water column TDN | 0.13 |
| Epilithon sites | Water quality groups | 0.71 |
| | Latitude | 0.68 |
| | Water column DIC | 0.66 |
| | Water column Temperature | 0.65 |
| | Sediment pH | 0.61 |
| | Longitude | 0.49 |
| | Water column Cations | 0.48 |
| | Slope | 0.45 |
| | NLCD 4 | 0.44 |
| | NLCD 5 | 0.32 |
| | Water column PAR | 0.31 |
| | Sediment Cations | 0.31 |
| | Water column Dissolved Oxygen | 0.26 |
| | Watershed area | 0.25 |
| | Water column Anions | 0.16 |

Table S3. Table listing environmental variables that were statistically significant in the NMDS (Figure 3 of the main manuscript). Variables are listed from highest $R^2$ to lowest, for both all substrate types (top) and epilithon sites only (bottom). DIC: dissolved inorganic carbon; NLCD: National Land Cover Dataset; PAR: Photosynthetically active radiation; TDN: total dissovle dnitrogen

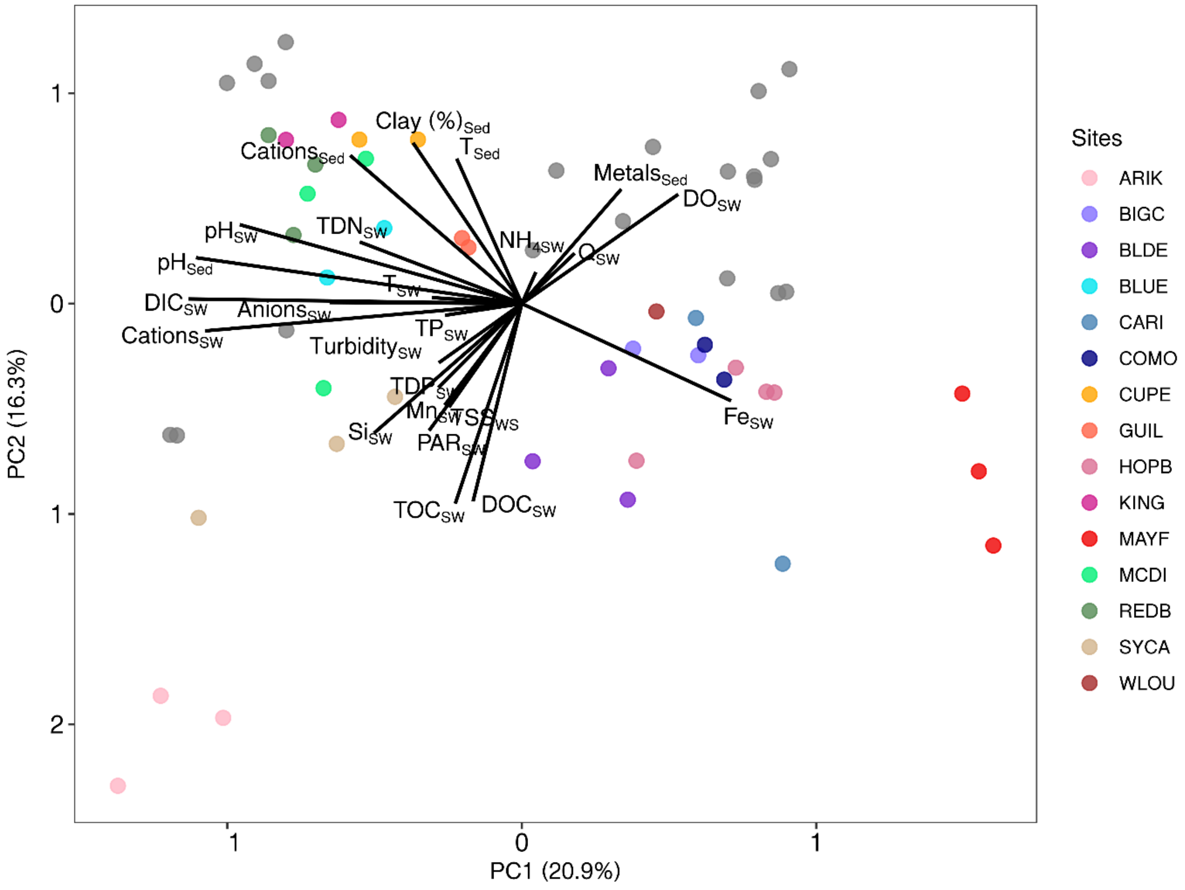


Figure S12. Principal component analysis (PCA) of environmental variables across NEON sites. Sed: sediment; SW: surface water; DIC: dissolved inorganic carbon; DO: dissolved oxygen; DOC: dissolved organic carbon; Fe: iron; Mn: manganese; NH4: ammonium; PAR: photosynthetically active radiation; Q: discharge; Si: silicate; T: temperature; TDN: total dissolved nitrogen; TDP: total dissolved phosphorus; TOC: total organic carbon; TP: total phosphorus.
